# Production of clinical grade patient iPSC-derived 3D retinal organoids containing transplantable photoreceptor cells

**DOI:** 10.1101/2025.03.04.640656

**Authors:** Laura R. Bohrer, Luke A. Wiley, Allison T. Wright, Bradley Hittle, Mallory J. Lang, Louisa M. Affatigato, Kimerly A. Powell, Lorena M. Haefeli, Ian C. Han, Robert F. Mullins, Edwin M. Stone, Budd A. Tucker

## Abstract

Neurodegenerative conditions that affect the retina are currently the leading cause of incurable blindness in the developed world. Although gene and drug therapies are being developed to slow disease progression in some cases, restorative cell replacement approaches are needed for patients with significant vision impairment due to retinal degeneration. While a variety of different cell types have been evaluated in the context of retinal cell replacement, induced pluripotent stem cells (iPSCs), which can be generated and delivered as an autologous therapeutic, are in many ways the most attractive donor cell source currently available. Like embryonic stem cells, iPSCs must be differentiated into the target therapeutic cell type prior to transplantation. For instance, for patients with retinitis pigmentosa who have primary photoreceptor cell disease, photoreceptor cell derivation and enrichment are required prior to transplantation. Although other effective retinal differentiation protocols exist, they are often not fully compatible with clinical manufacturing. In this study, we report development of a xeno-free 3D retinal differentiation protocol based on the most robust adherent/non-adherent 3D differentiation strategies published to date. In addition, we demonstrate that while iPSC reprogramming efficiency is enhanced under reduced oxygen tension (i.e., 5%), efficient embryoid body and subsequent retinal organoid production require standard oxygen levels (i.e., 21%). Finally, we show that photoreceptor precursor cells obtained from 3D retinal organoids derived using the developed protocol under current good manufacturing practices (cGMP) survive in the subretinal space of dystrophic *Pde6b*-null rats for 1-month post-transplantation and form new synaptic connections with host bipolar neurons.

**Significance statement:** In this study, we report development of a xeno-free, cGMP compliant iPSC-3D retinal differentiation protocol for production of transplantable photoreceptor precursor cells.

## Introduction

Neurodegenerative disorders, ranging from relatively common conditions such as Parkinson’s disease to rare forms of inherited retinal degeneration, involve death of primary neurons of the central nervous system, resulting in severe cognitive, neurologic, and/or visual impairment. As CNS neurons have limited regenerative capacity, current drug- and gene-therapies for these conditions are primary designed to prevent further neuronal cell death. For patients with advanced disease, cell replacement will be needed to restore lost cells and regain function. In the eye, loss of the light-sensing photoreceptor cells occurs in a wide variety of conditions including age-related macular degeneration and retinitis pigmentosa (RP). Progressive loss of photoreceptor cells results in progressive vision loss and significant impact on daily activities including reading and driving. As neurons of the inner retina (including bipolar and retinal ganglion cells (1) persist for years following photoreceptor cell death, there is hope that photoreceptor cell replacement will restore vision and enable patients to regain their normal activity. As the retinal environment for transplantation is favorable (e.g., the retina is devoid of myelin and associated growth inhibitory proteins), stem cell-based photoreceptor cell replacement is one of the most promising therapeutic strategies currently being developed.

Over the past two decades, a variety of different donor cell types have been evaluated in the context of therapeutic photoreceptor cell replacement. These range from embryonic and induced pluripotent stem cells (iPSCs), to fate restricted mesenchymal stem cells (MSCs), retinal progenitor cells, and photoreceptor precursor cells. The ability of iPSCs to be generated from the patient in need has contributed to the widespread popularity of this technology across nearly all fields of neural engineering. Extensive data from the field of transplant biology clearly demonstrate that immunologically matched tissues are superior to unmatched tissues when graft function and longevity are considered (2–4). Furthermore, patient derived iPSCs avoid ethical and logistic challenges that are often associated with the use of primary fetal cell types (e.g., retinal progenitor and photoreceptor precursor cells). That said, one of the short comings of iPSC technology is that, like embryonic stem cells, they must be differentiated into the target cell type prior to transplantation (i.e., transplantation of undifferentiated pluripotent stem cells would likely result in teratoma formation). While excellent retinal differentiation protocols have been reported, they often rely on the use of reagents derived from non-human sources. For instance, extracellular matrix molecules such as Matrigel and cell culture reagents such as Fetal Bovine Serum (FBS) are common in state-of-the-art differentiation paradigms. To address this concern, in 2017 we published a modified version of a 3D retinal differentiation protocol initially reported by Eiraku and Sasai (5) that relied exclusively on the use of validated xeno-free components (6). While effective, organoids generated using this approach contain both retinal and non-retinal forebrain tissues. Although retinal tissue could be mechanically dissected and grown free from non-retinal tissues, this process was difficult, inefficient, and introduced additional opportunity for microbial contamination. In addition, retinal organoids derived using our previously published protocol generally lacked the well laminated structure normally present in the neural retina. Newer 3D differentiation protocols that utilize a combined adherent/non-adherent culture strategy have enabled efficient production of laminated retinal organoids free from contaminating non-retinal cell types (7, 8). This is important considering that we have recently demonstrated how normal retinal lamination (i.e., an outer photoreceptor cell layer separated from an inner layer of bipolar, muller, amacrine and horizontal cells) can be exploited to enable isolation of an enriched photoreceptor cell population for downstream cell replacement applications (9).

In this study, we report development of a xeno-free 3D retinal differentiation protocol based on the most robust adherent/non-adherent 3D differentiation strategies published to date (8, 10–12). We demonstrate that while iPSC reprogramming efficiency is enhanced under low oxygen tension, efficient embryoid body and subsequent retinal organoid production require standard oxygen levels (i.e., ∼21%). We subsequently demonstrate that xeno products such as dispase, Matrigel and FBS can be replaced with xeno-free reagents such as ReLeSR, CELLstart and KnockOut Serum Replacement (KSR), respectively. Following subretinal transplantation, photoreceptor precursor cells obtained from 3D retinal organoids derived under current good manufacturing practices (cGMP) survive in the subretinal space of dystrophic *Pde6b*-null rats and form new synaptic connections with host bipolar neurons.

## Materials and Methods

### Ethics statement

This study was approved by the Institutional Review Board of the University of Iowa (project approval #200202022) and adhered to the tenets set forth in the Declaration of Helsinki, with patients providing written, informed consent. All rat experiments were conducted with the approval of the University of Iowa Animal Care and Use Committee (Animal welfare assurance #1031317) and were consistent with the ARVO Statement for the Use of Animals in Ophthalmic and Vision Research.

### Production of patient-derived iPSCs

Reprogramming and clonal expansion were performed using the Cell X™ platform (Cell X Technologies Inc), which is contained within a cGMP compliant Biospherix Xvivo system (BioSpherix, Ltd.) (10). Briefly, dermal fibroblasts isolated from 3 mm dermal punch biopsies obtained from the individuals listed in **Table 1** were cultured at 37° C, 5% CO2, and 20% O2 and transduced using the CytoTune 2.0 kit (Thermo Fisher Scientific; Cat#A16517), a non-integrating Sendai viral reprogramming kit (6) at 20% O2. After 5 days, cells were passaged onto rhlaminin 521 (5 μg/ml; Biolamina; Cat#LN521-02) coated dishes. To enhance reprogramming efficiency, the oxygen tension was reduced from 20% to 5% at 1 week following transduction. At passage 10, iPSC lines were subject to karyotyping and scorecard analysis as described previously (10).

### RNA isolation and hPSC ScoreCard™ analysis

For TaqMan hPSC ScoreCard™ analysis, total RNA was isolated using the NucleoSpin RNA purification kit (Macherey-Nagel; Cat#740955). cDNA was generated from 1 μg of RNA using VILO cDNA synthesis kit (Thermo Fisher Scientific; Cat#11754050). cDNA was added to a TaqMan hPSC scorecard plate (Thermo Fisher Scientific; Cat#A15870) and amplified using a QuantStudio 6 Flex real-time PCR system. Results were analyzed using Thermo Fisher’s cloud-based analysis suite.

### Xeno-free derivation of iPSC-derived retinal organoids

Retinal differentiation was performed with modifications for GMP-compliance (8, 10, 11). Briefly, iPSCs were cultured on recombinant human rhlaminin 521 coated cell culture plates in E8 medium (Thermo Fisher Scientific; Cat#A1517001). To generate embryoid bodies (EBs), iPSCs were lifted using ReLeSR™ (STEMCELL Technologies; Cat#05872) rather than dispase (as per our standard protocol (13). Cell clusters were transitioned from E8 to neural induction medium (NIM-DMEM/F12 (1:1) (Thermo Fisher Scientific; Cat#11320033), 1% N2 supplement (Thermo Fisher Scientific; Cat#17502048), 1% non-essential amino acids (Thermo Fisher Scientific; Cat#11140050), 1% Glutamax (Thermo Fisher Scientific; Cat#35050061), 2 μg/mL heparin (MilliporeSigma; Cat#H3149) and 0.2% Primocin (InvivoGen; Cat#ant-pm-2)) over a four-day period. On day 6, NIM was supplemented with 1.5 nM rhBMP4 (R&D Systems; Cat#314-BP-05/CF). On day 7, EBs were adhered to Matrigel™ (Corning; Cat#354234 – standard protocol) or the following xeno-free substrates: CELLstart™ (Thermo Fisher Scientific; Cat#A1014201), a mixture of recombinant human laminin 111 (LN111) (5 μg/ml; Biolamina; Cat#LN111-02), recombinant human nidogen-1 (5 μg/ml; R&D Systems; Cat#2570-ND-050) and recombinant human collagen IV (18 μg/ml; MilliporeSigma; Cat#CC076), or recombinant human LN111 only. BMP4 was gradually transitioned out of the NIM over seven days. On day 16, the media was changed to retinal differentiation medium (RDM - DMEM/F12 (3:1), 2% B27 supplement (Thermo Fisher Scientific; Cat#17504044), 1% non-essential amino acids, 1% Glutamax and 0.2% Primocin). On day 30 the entire outgrowth was mechanically lifted using the back end of a 1 ml pipette tip and transferred to ultra-low attachment flasks in xeno-free 3D-RDM (RDM plus 10% Fetal Bovine Serum (FBS; Atlas Biologicals; Cat#F-0500-A) (as per our standard protocol) or 10% KnockOut™ Serum Replacement (KSR; Thermo Fisher Scientific; Cat#10828028), 100 µM taurine (MilliporeSigma; Cat#T0625), 1:1000 chemically defined lipid concentrate (Thermo Fisher Scientific; Cat#11905031), and 1 µM all-trans retinoic acid (MilliporeSigma; Cat#R2625). The cells were fed three times per week with 3D-RDM until differentiation day 100 when media was switched to 3D-RDM minus (i.e., 3D-RDM without all-trans retinoic acid). Organoids were feed 3 days per week with 3D-RDM minus until harvest.

### Dissociation of retinal organoids

Day 160 retinal organoids were allowed to settle and washed with EBSS buffer (Worthington Biochemical Corporation; Cat#LS006361). Organoids were dissociated with 35 units/mL of papain (Worthington Biochemical Corporation; Cat#LK003176) and 150 units/mL DNase I (Worthington Biochemical Corporation; LS006361) in EBSS buffer. Organoids were incubated in a 37°C water bath for 45 minutes and were pipetted every 15 minutes to break up remaining tissue chunks. Following dissociation, cells were put through a 40 μm strainer, pelleted and resuspended in HBSS plus calcium, magnesium and N-acetyl-cysteine (NAC, 0.5 mM) (Spectrum Chemical Corporation; Cat#AC800).

### Animals

We recently described the generation of a *Pde6b-*null rat model of retinal degeneration via CRISPR-mediated genome editing (14). This model has an early onset rapidly progressive disease resulting in near complete loss of photoreceptor cells by two-months of age. For the immunosuppression protocol development studies, 2-month-old *Pde6b*-deficient rats (N=6, 3M, 3F) were used. For the subsequent cell replacement studies two groups of 2-month-old *Pde6b*-deficient rats were used. Group 1 was sacrificed at 3-days (N=12; 6M, 6F) and group 2 was sacrificed at 4-weeks (N=12; 6M, 6F) following subretinal injection of 2,000,000 human iPSC-derived retinal cells. In each experiment, only one eye received subretinal injections and the contralateral eye was used as a control.

### Immunosuppression and subretinal injection of retinal cell suspension

For immunosuppression prior to xenograft injections, rats were first treated with intraperitoneal injection of cyclosporine (10 mg/kg) daily for seven days starting two days prior to subretinal injection of human iPSC-derived retinal cells (i.e., day −2). Following cyclosporine injection, immunosuppression was continued via cyclosporine-supplemented drinking water (100 μg/mL) ad libitum until animal euthanasia.

For subretinal injections of retinal cell suspensions, rats were placed under anesthesia with a weight-dependent cocktail of intraperitoneal ketamine (91 mg/kg) and xylazine (9.1 mg/kg). Once anesthetized, eyes were dilated with 1% tropicamide. Using an operating microscope, a limited conjunctival peritomy was created in the temporal quadrant using 0.12 mm forceps and Vannas scissors. Subretinal injections were performed using a 33-gauge blunt-tipped Hamilton syringe as previously described (14–17). All injections were performed by a fellowship-trained vitreoretinal surgeon (I.C.H.).

Each eye was immediately imaged post-injection using a rodent-specific fundus camera and optical coherence tomography instrument (Micron IV+OCT; Phoenix MICRON Image-Guided OCT2, Phoenix Laboratories) to confirm the presence and location of a subretinal bleb and to check for the absence of complications (e.g., perforation of the retina into the vitreous cavity or subretinal hemorrhage) as described previously (14–17). The contralateral eye of each animal was used as a no injection control.

### Organoid and tissue processing for histological analysis

Retinal organoids were fixed with 4% paraformaldehyde for 30-60 minutes at room temperature and equilibrated to 15% sucrose in PBS, followed by 30% sucrose. Organoids were cryopreserved in 50:50 solution of 30% sucrose/PBS: tissue freezing medium (Electron Microscopy Sciences; Cat#72592) and cryosectioned (15 μm).

For eyes that received subretinal injection of a single cell suspension, the injection site was identified by fundus examination, and a limbal suture was placed at the corresponding clock hour of the injection for orientation when embedding tissues for sectioning, as described previously (14–17). Rat eyes were enucleated and fixed in 4% paraformaldehyde for at least 1 hour prior to dissection of the anterior segment. The posterior eye cup was fixed in 4% paraformaldehyde overnight at 4°C and rinsed in increasing concentrations of sucrose. The eye cup was oriented using the previously-placed limbal suture to identify the injection site, embedded in 2:1 solution of Tissue-Tek OCT compound (VWR International; Cat#4583) and 20% sucrose, flash-frozen in liquid nitrogen, stored at −80°C, and sectioned at a thickness of 7 µm using a Microm HM505E (Microm) or Leica cryostat as previously described (14–17).

### Immunohistochemistry and confocal microscopy

Retinal organoid sections were blocked in PBS containing 5% normal donkey serum (MilliporeSigma; Cat#S30), 3% bovine serum albumin (BSA; Research Products International Corp.; Cat#A30075), and 0.1% triton-x. Retinal sections were blocked in PBS containing 3% BSA at room temperature for an hour. Primary antibodies (**Table S1**) were diluted in blocking buffer and allowed to incubate for 2 hours at room temperature or overnight at 4°C. Following washes in PBS, sections were incubated with appropriate Alexa Fluor fluorescent-conjugated secondary antibodies (**Table S1**). 4’,6-Diamidino-2-phenylindole dihydrochloride (DAPI; Thermo Fisher Scientific; Cat#62248) was used to counterstain cell nuclei. Sections were mounted using Aqua-Mount Mounting Medium (Thermo Fisher Scientific; Cat#14-390-5) and then visualized using a Leica TCS SPE upright confocal microscope system (Leica Microsystems).

## Results

### Replacement of xeno products in retinal differentiation protocol

Although inclusion of animal-derived products (such as Matrigel and FBS) in therapeutic stem cell differentiation pipelines is not prohibited by regulatory authorities, use of xeno-free components is strongly encouraged. In addition to the increased risk of containing pathogenic contaminants, animal-derived products often suffer from significant lot-to-lot variability. Inconsistency of supplements used in the manufacturing process can have a major impact on protocol reproducibility and potency of the therapeutic product. As depicted in **Figure 1A**, the most robust retinal differentiation protocol that we have used to date is based on that first published by Meyer et. al. (7), which uses a series of adherent and non-adherent cell culture steps to produce free-floating retinal organoids. This protocol begins with passage of adherent iPSCs from LN521 coated tissue culture plates using ReLeSR into ultra-low adhesion dishes in 75% Essential 8 (E8) media and 25% Neural Induction Media (NIM) (**Figure 1A**).

**Figure 1.**
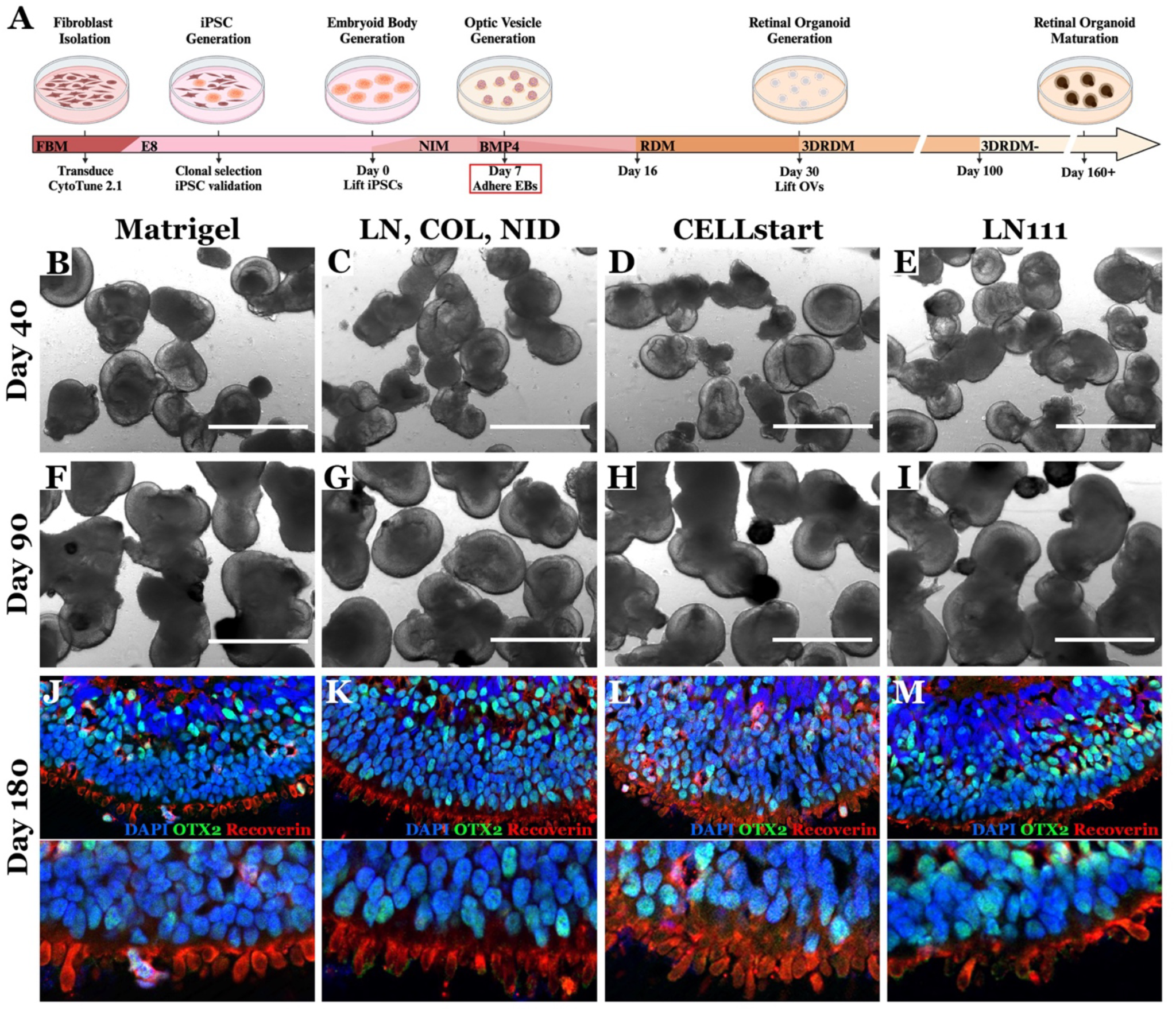
Replacement of Matrigel with xeno-free products designed to promote embryoid body attachment and outgrowth. A) Schematic of our previously reported retinal differentiation protocol highlighting the animal derived reagent needing to be replaced (i.e., Matrigel coated dishes are used to adhere embryoid bodies at differentiation day 7). At differentiation day 7, embryoid bodies were attached to cell culture surfaces coated with 1 of 4 different substrates: Matrigel (B, F, J); or recombinant human laminin 111 (LN), Collagen IV (COL), and nidogen (NID) (C, G, K); CELLstart (D, H, L); recombinant human laminin 111 (LN111) (E, I, M). Representative phase micrograph images of early retinal organoids at differentiation days 40 (B-E) and 90 (F-I). Immunohistochemical staining of mature retinal organoids at differentiation day 180 using antibodies targeted against the photoreceptor cell markers OTX2 (green) and recoverin (red). DAPI (blue) was used as a nuclear counterstain. Scale bar= 1 mm.

Cultures are fed with 50% E8 and 50% NIM on differentiation day 1, 25% E8 and 75% NIM on differentiation day 2 and 100% NIM on differentiation day 3. On differentiation day 6, NIM is supplemented with BMP4 to bias retinal cell fate specification (8, 18) (**Figure 1A**). In the standard non-xenofree protocol on differentiation day 7, resulting embryoid bodies are plated onto Matrigel coated cell culture surfaces to enable attachment, outgrowth of neural progenitor cells, and formation of optic vesicle-like structures (**Figure 1A**). Matrigel, which is derived from the mouse chondrosarcoma Engelbreth-Holm-Swarm cell line, is particularly prone to lot-to-lot variability (e.g., lot 11524001 = 11.1 mg/ml total protein and lot 27624003 = 7.9 mg/ml total protein). To identify a suitable Matrigel replacement for embryoid body attachment, on differentiation day 7, embryoid bodies were plated onto dishes coated with 1 of 4 different substrates (**Figure 1B, F, & J** = Matrigel; **Figure 1C, G & K** = recombinant human laminin 111, collagen IV, and nidogen-1; **Figure 1D, H, & L** = CELLstart; and **Figure 1 E, I, & M** = recombinant human laminin 111). BMP4 was gradually transitioned out of the NIM over seven days (100% on Day 6, 50% on day 9, 25% on day 12, 12.5% on day 14). On differentiation day 16, NIM was switched to BMP4-free Retinal Differentiation Media (RDM). At differentiation day 30, cultures were mechanically lifted using the back end of a 1 ml pipette tip and cultured in 3DRDM in suspension until differentiation day 100, at which time the media is switched to 3DRDM-(no retinoic acid). No difference in early neural epithelial layer development (**Figure 1B-E**) or retinal lamination (**Figure 1F-I**) were detected by phase microscopy between any of the substrates used for embryoid body attachment. Likewise, the number of organoids present in each condition was unaffected by plating substrate. The impact of embryoid body plating substrate on retinal organoid maturation was evaluated at differentiation day 180 via immunohistochemical analysis using primary antibodies targeting OTX2 and recoverin (**Figure 1J-M**). Again, we did not detect a significant difference between retinal organoids generated from embryoid bodies plated on any of the xeno-free substrates as compared to those cultured on Matrigel. All organoids developed clearly demarcated inner and outer nuclear layers (**Figure 1J-M**). Furthermore, the outer nuclear layer of all retinal organoids contained OTX2 and recoverin positive photoreceptor cells that extended outer segment-like structures (**Figure 1J-M**).

As indicated above, optic vesicle-like structures are lifted at differentiation day 30, during which time the cell culture media is switched from RDM to 3DRDM (**Figure 2A**). Unlike RDM, 3DRDM contains fetal bovine serum (FBS), which is added to improve long term cell survival and retinal organoid maturation (**Figure 2A**).

**Figure 2.**
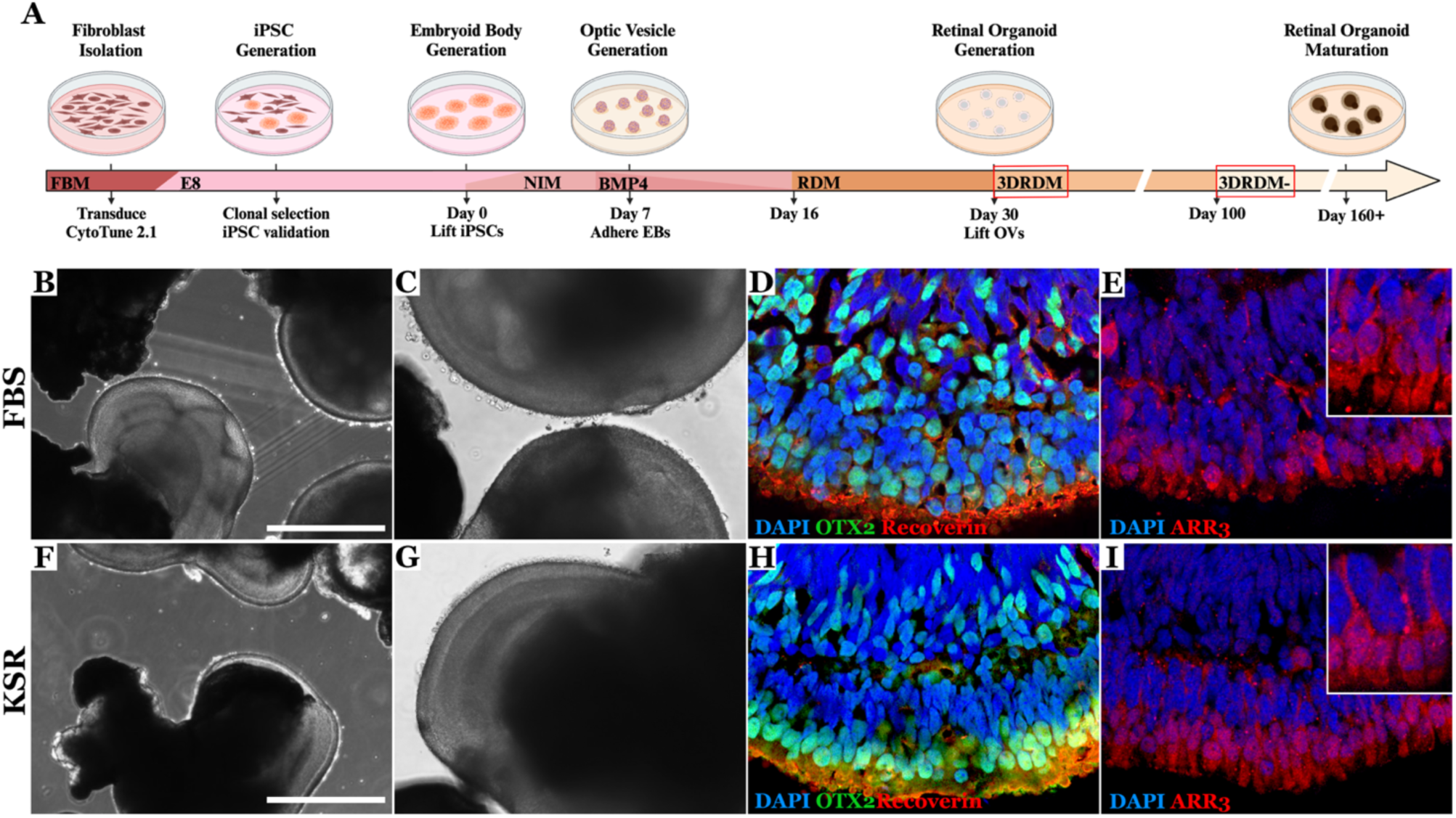
Replacement of FBS with xeno-free knockout serum replacement. **A)** Schematic of our previously reported retinal differentiation protocol highlighting the animal derived reagent needing to be replaced (i.e., FBS is added to 3DRDM beginning at differentiation day 30). **B-I)** Phase (B-C & F-G) and immunohistochemical analysis of retinal organoids at differentiation day 160 that were matured in either FBS (B-E) or KSR (F-I). For immunohistochemical analysis antibodies targeted against OTX2, recoverin, and ARR3 were used (nuclei were counterstained using DAPI). Scale bar= 400 um.

One of the most well characterized xeno-free FBS alternatives that is currently available as a clinical grade reagent is knockout serum replacement (KSR). To determine if KSR could be used in place of FBS, we differentiated iPSCs and used CELLstart in place of Matrigel for embryoid body attachment. At differentiation day 30, half of the organoids were placed in 3DRDM supplemented with FBS and half were placed in 3DRDM supplemented with KSR. Retinal organoids were assessed at differentiation day 160. Those grown in FBS (**Figure 2B-E**) and KSR (**Figure 2F-I**) had similar morphology with clear lamination and extension of outer segment like projections. Immunohistochemical analysis revealed expression of the photoreceptor cell markers OTX2 and Recoverin (**Figure 2D, H**) as well as the cone phototransduction protein ARR3 (**Figure 2 E, I**).

### Influence of oxygen tension on retinal differentiation capacity

Although cellular reprogramming has been well characterized, the process of converting human dermal fibroblasts to iPSCs following forced expression of OCT4, SOX2, KLF4 and c-MYC is often inefficient. Many strategies designed to enhance reprogramming efficiency have been evaluated, the vast majority of which rely on addition of small molecule inhibitors (19, 20) or modification of the reprogramming factor cocktail (21). In 2009, Yoshida and colleagues first demonstrated that culturing human dermal fibroblasts under 5% oxygen for 1, 2, or 3 weeks, beginning 1-week post-transgene expression, enhanced reprogramming efficiency by as much as 4-fold (22). As modifying oxygen tension within the cell culture incubator does not require additional cGMP compliant reagents, we have incorporated hypoxia into our cGMP iPSC generation pipeline (10). For donors such as the one presented in **Figure 3**, reprogramming under reduced oxygen tension is essential. To demonstrate, dermal fibroblasts were passaged 5-days following viral transduction and split at an equal density into two separate LN521 coated cell culture dishes (**Figure 3A & B**). At 7 days post-transduction one plate of cells was placed in an incubator maintained at 5% oxygen tension and the other plate was placed in an incubator maintained at 20% oxygen tension. The number of iPSC colonies present two weeks later was determined to be approximately 30-fold greater for cultures maintained at 5% oxygen tension versus those maintained at 20% oxygen tension (**Figure 3C, E, F & G** – colony analysis was performed using images collected via the CellX device and an automatic iPSC detection algorithm as we have previously described (23)). Interestingly, iPSC colonies generated under 5% oxygen tension were larger in diameter and as such ready to pick sooner than those generated under 20% oxygen tension (**Figure 3E-G**).

**Figure 3.**
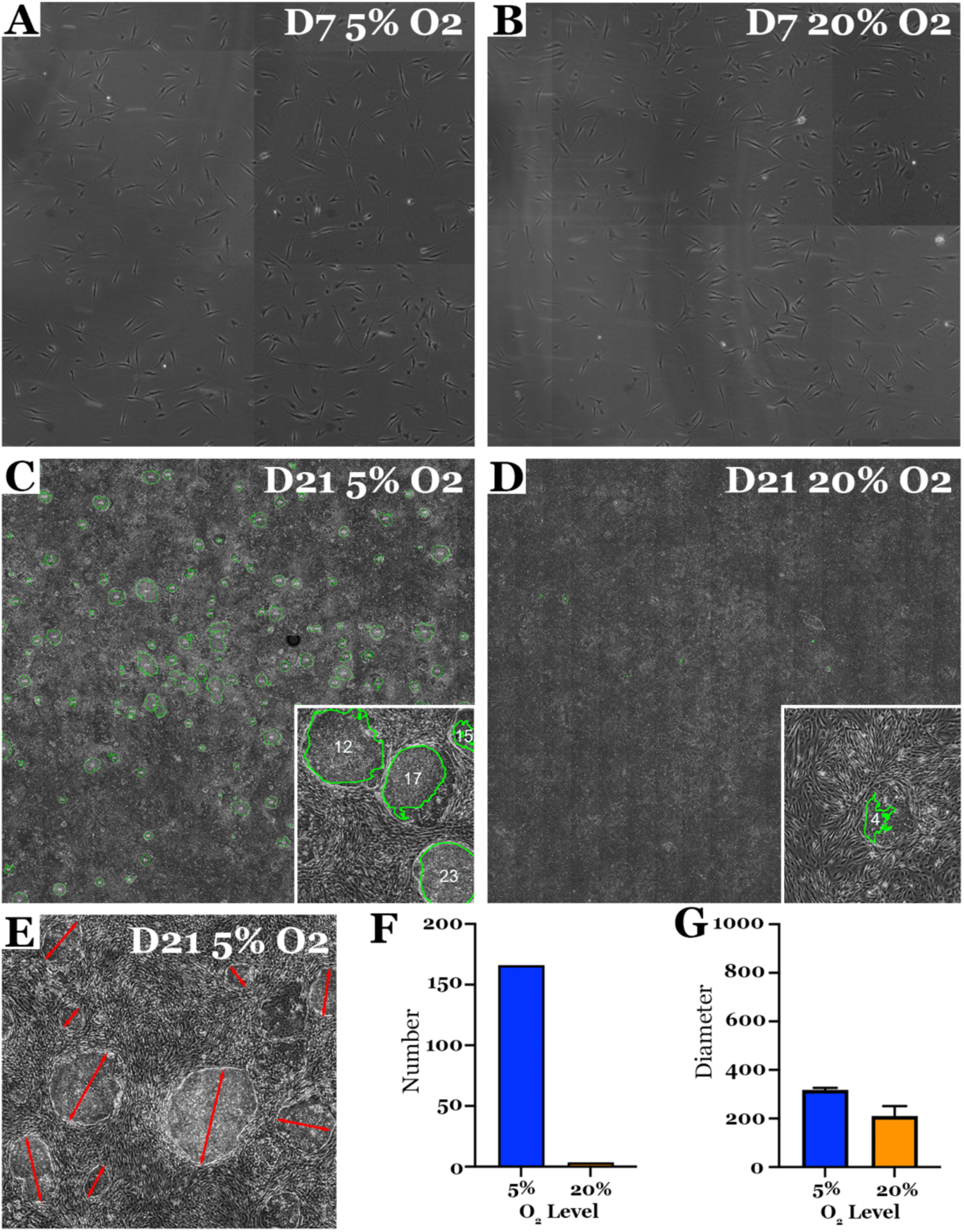
Reduced oxygen tension enhances reprograming efficiency of difficult to reprogram patient lines. **A-D)** Patient derived dermal fibroblasts cultured at 20% oxygen tension were transduced with sendai viral vectors driving expression of OCT4, SOX2, KLF4 and C-MYC (CytoTune2). At 5 days post-transduction cells were passaged at a density of ∼2,000 cells per cm^2^ into two separate LN521 coated 10cm cell culture dishes. At 7 days post-transduction one plate of cells was cultured at 5% oxygen tension and the other was cultured at 20% oxygen tension. Plates were imaged every other day using Cell X platform beginning on day 7 (A - 5% O2; B - 20% O2) and continued through day 21 (C - 5% O2; D - 20% O2). **E-G)** The number (F) and diameter (E, G) of iPSC colonies present in the entire plate was determined using automated iPSC colony detection and segmentation software.

To evaluate the impact of oxygen tension on retinal differentiation, iPSCs lines generated at 5% oxygen were split and either maintained at 5% oxygen or cultured under 20% oxygen for 2 passages prior to initiating differentiation. Cells were differentiated using the newly developed xeno-free differentiation protocol described above. As compared to iPSCs maintained at 5% oxygen, cells maintained at 20% oxygen were contained in densely packed colonies, had a high nucleus to cytoplasm ratio, lacked phase bright edges and had increased self-renewal scores as assessed by the TaqMan hPSC ScoreCard analysis (**Figure 4B, G & L**). In addition, as determined by RNA abundance, at 7 days post-differentiation there were significantly more embryoid bodies present in iPSC cultures maintained under 20% oxygen tension than those cultured under 5% oxygen tension (**Figure 4C, H & N**).

**Figure 4.**
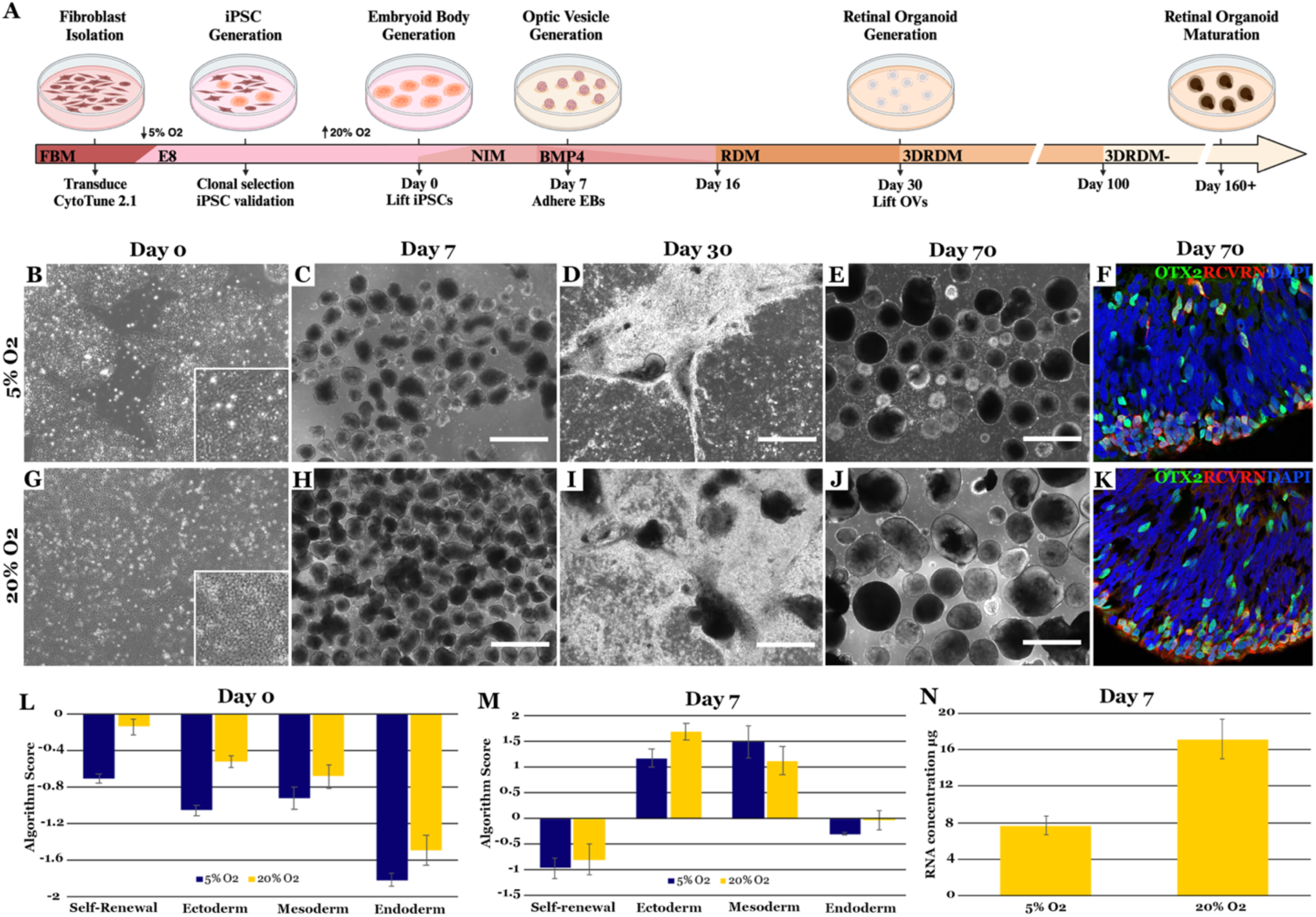
Retinal differentiation of iPSCs at 5 or 20% oxygen. **A)** Schematic of the cGMP-compliant retinal differentiation protocol. **B-K:** Representative micrographs of day 0 iPSCs (B, G), day 7 EBs (C, H), day 30 attached optic vesicles (D, I) and day 70 lifted retinal organoids (E, J, F & K). Immunohistochemical staining of retinal organoids at day 70 (F, K) for the photoreceptor cell markers OTX2 (green) and Recoverin (red). DAPI (blue) was used as a nuclear counterstain. Scale bar=1 mm. **L-M)** TaqMan hPSC Scorecard analysis comparing iPSCs (L) and day 7 embryoid bodies (M) at 5% or 20% oxygen. **N)** Concentration of RNA isolated from day 7 embryoid bodies generated and cultured at 5% or 20% oxygen.

Interestingly, there was no significant difference between lines maintained in 5% oxygen versus 20% oxygen when expression of germ layer specific genes was evaluated, suggesting that oxygen tension alters embryoid formation potential rather than cell fate decision (**Figure 4M**). By 30 days of differentiation, optic vesicles were identified in both 5% and 20% oxygen conditions. Interestingly, as with embryoid body formation, there was significantly fewer optic vesicles present in cultures maintained at 5% oxygen than those maintained at 20% oxygen (**Figure 4D & J**). Since iPSCs differentiated under 20% oxygen generated significantly more embryoid bodies and in turn optic vesicles that go on to develop into mature retinal organoids (**Figure 4E, F, J & K**), all subsequent studies were done at this oxygen level.

### Subretinal transplantation of xeno-free iPSC-derived retinal cells

To test the utility of our amended xeno-free retinal differentiation protocol for derivation of transplantable photoreceptor cells, subretinal injections were performed in the *Pde6b*-deficient rat model (14). The *Pde6b*-deficient rat has a rapidly progressive retinal degeneration with near complete loss of photoreceptor cells by 2-months of age (14), which makes it ideal for evaluation of post-transplant donor cell integration (i.e., the lack of host photoreceptor cells at the time of transplantation mitigates concerns pertaining to donor cell antigen transfer (24–26)). As previously reported in other rodent models of retinal degeneration (27–32), disease induced activation of retinal microglia can be detected as early as post-natal day 14 in *Pde6b-/-* rats (**Supplemental Figure 1**). In addition to expressing IBA1 and being rounded in shape compared to their normal ramified appearance in the non-diseased state, these cells also express CD68, a lysosomal marker typically present in activated, phagocytosing microglia (**Supplemental Figure 1**). By 28-days of age, activated microglia are still present within the degenerating photoreceptor cell layer and subretinal space of the *Pde6b*-/- rat (**Supplemental Figure 1**). To determine if the systemic immunosuppression protocol described in the methods section mitigates immune cell infiltration and xenograft rejection, 500,000 iPSC derived retinal cells were injected into the subretinal space of immunosuppressed versus naïve animals. Compared to non-immunosuppressed controls, cyclosporine treated animals had an appreciable decrease in the number of Iba1-postive cells in the neural retina (**Supplemental Figure 2B-E**). Importantly, no influx of CD45 positive infiltrating monocytes or other leukocytes was observed from cyclosporine-treated animals (**Supplemental Figure 2F-I**).

Following confirmation of immune suppression, xeno-free iPSC-derived retinal organoids were dissociated, and 2 million cells were injected into the subretinal space of one eye. Immunohistochemical analysis was performed at 3- and 30-days post-injection. To visualize human iPSC-derived cells, the human-specific antibodies human nuclear antigen (HNA), Stem101 (AKA Ku80), Stem121, and human mitochondrial antigen (HMA) were evaluated (**Supplemental Figure 3**). As HNA was found to be the most specific with the least amount of background labeling, this antibody was used in subsequent experiments.

At 3-days post-transplantation, HNA-positive cells were present in a continuous, multi-layered sheet within the subretinal space immediately under the degenerative host inner nuclear layer (**Figure 5A-E**), demonstrating successful transplantation. HNA-positive cells expressed the photoreceptor cell marker recoverin (**Figure 5B** - expressed in both rods and cones) and the photoreceptor transcription factor OTX2 (**Figure 5C** – this marker is also expressed in mature bipolar cells). HNA positive cells did not express the retinal ganglion/amacrine cell marker Calretinin (**Figure 5D**) or the bipolar cell marker PKCα (**Figure 5E**). There were pigmented RPE cells in the injected area positive for CRALBP (**Figure 5F**) that were easily identified by H&E staining (**Figure 5G**).

**Figure 5.**
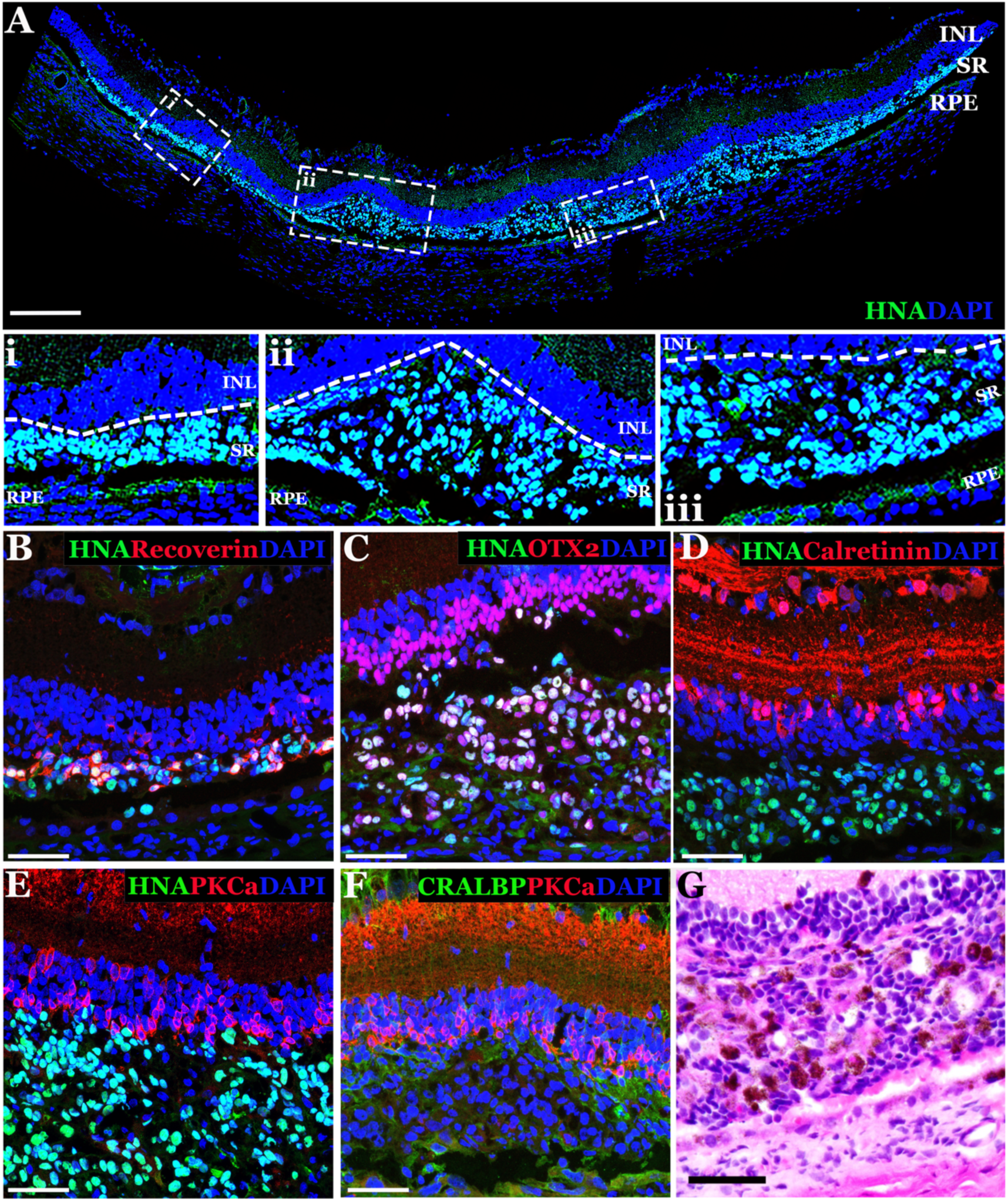
Xenotransplanted human iPSC-derived retinal cells take residence in the subretinal space of immunosuppressed *Pde6b* KO rats at 3-days post-injection. **A-F)** Immunohistochemical analysis of rat retinas at 3 days following subretinal injection of clinical grade human iPSC-derived retinal cells. Antibodies targeting the human nuclear antigen (HNA) (A-E, used to identify donor cells of human origin), recoverin (B - photoreceptor cell marker), OTX2 (C - photoreceptor and bipolar cell marker), calretinin (D - ganglion/amacrine cell marker), PKCα (E & F - bipolar cell marker) and CRALBP (F - RPE cell marker) were used. Representative panoramic image displaying the extent of the retinal transplant (A). Dashed white boxes demarcate areas of higher magnification displayed below the panoramic (i, ii, and iii). **G)** H&E staining demonstrating presence of pigmented donor RPE cells present in the graft. DAPI (blue) was used as a nuclear counterstain. Abbreviations for retinal layers in A: Inner Nuclear Layer (INL), subretinal space (SR), Retinal Pigmented Epithelium (RPE). Scale bars = 200 μm (A) and 50 μm (B-G).

At 30 days post-transplant, host photoreceptor cells were largely absent (**Figure 6A**), but recoverin positive photoreceptor cells were present almost exclusively within the transplant region (**Figure 6B-D’**). While fewer HNA positive donor cells remained at 30 days post-transplantation, the vast majority expressed the photoreceptor cell marker recoverin (**Figure 6B**). Similarly, s-opsin positive cones were almost exclusively HNA positive (**Figure 6C**). Donor HNA positive iPSC-derived photoreceptor cells remained at the outer limits of the neural retina juxtaposed to PKCα positive bipolar cells (**Figure 6D-D’**). Staining for the synaptic marker synaptophysin revealed expression between recoverin positive photoreceptor cells and host bipolar neurons only (**Figure 6D-D’**), suggesting that transplanted cells are attempting to form new synaptic connections with the residual host retina.

**Figure 6.**
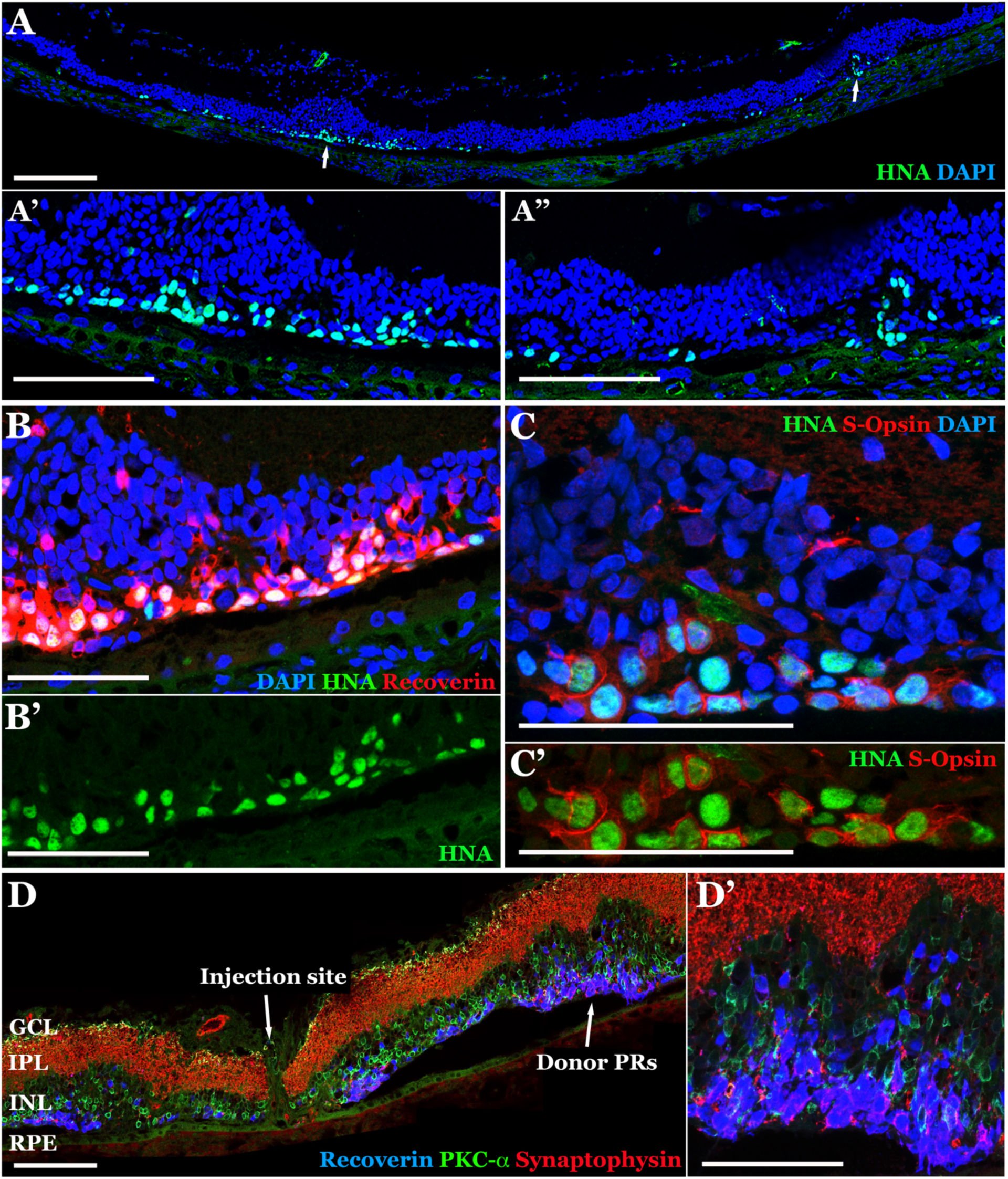
Xenotransplanted human iPSC-derived photoreceptor precursor cells survive within the subretinal space of host degenerative rat retina at 30-days post-transplantation. **A-D)** Immunohistochemical analysis of rat retinas at 30 days following subretinal injection of clinical grade human iPSC-derived retinal cells into *Pde6b*-deficient animals. Antibodies targeting the human nuclear antigen (HNA) (A-C, used to identify donor cells of human origin), recoverin (B & D - photoreceptor cell marker), s-Opsin (C – cone photoreceptor cell marker), PKCα (E & F - bipolar cell marker), and synaptophysin (D – synaptic marker) were used. Abbreviations for retinal layers in D: Ganglion Cell Layer (GCL), Inner Plexiform Layer (IPL), Inner Nuclear Layer (INL) and Retinal Pigmented Epithelium (RPE). Scale bars = 200 μm (A & D) and 100 μm (A’-C’ & D’).

## Discussion

There remains a great clinical need for stem cell mediated photoreceptor cell replacement therapy with possible application in a wide variety of retinal degenerative diseases. Although such strategies have enormous potential, doubt was cast on the validity of early photoreceptor cell replacement studies due to the discovery that material including proteins such as GFP could be transferred between donor and host photoreceptor cells following subretinal transplantation (25, 26, 33), thus temporarily dampening enthusiasm. Fortunately, recent experiments performed in animal models of end stage disease have reestablished belief in the therapeutic potential of pluripotent stem cell mediated photoreceptor cell replacement. For instance, several groups have demonstrated that stem cell-derived photoreceptor cells transplanted into the subretinal space of rodents with advanced disease are able to form new synaptic connections with host inner retinal neurons and mediate some degree of functional recovery (34–36). In a recent study using modern methods to confirm donor cell identity, Gasparini and colleagues convincingly demonstrate functional integration of iPSC-derived cone photoreceptor cells and structural maturation marked by normal cellular polarization and reformation of an outer limiting membrane (37). These robust studies of photoreceptor cell replacement in translational models have further matured the field toward human clinical trials, emphasizing the need for efficient and reliable protocols meeting cGMP requirements for clinical-grade manufacturing.

To date, protocols used for photoreceptor cell derivation have typically relied on the use of reagents that are poorly compatible with clinical manufacturing. For instance, animal derived reagents such as dispase, Matrigel, and FBS are common in state-of-the-art retinal organoid production protocols. While we and others have reported xeno-free retinal differentiation protocols in the past, they have required manual picking and/or dissection of the desired retinal structures, which can be technically challenging, decreases throughput, and increases contamination risk (38–41). In addition, the absence of FBS in long term cultures often leads to increased cell death resulting in a loss of retinal lamination and neural rosette formation.

The primary goal of this study was to modify our previously published retinal differentiation protocol (42) to enable production of transplantable cells under current good manufacturing practices. In addition to removing animal derived products, we also evaluated the impact of low oxygen tension, which we use to enhance iPSC reprogramming efficiency. All procedures were performed in a cGMP compliant ISO class 5 Biospherix Xvivo unit to enable direct translation of the procedures to our clinical manufacturing suite. Unlike our previously reported cGMP compliant protocol, which was based on that of Nakano et. al. (43), our newly developed cGMP compatible differentiation paradigm leverages a base protocol first described and subsequently refined by Meyer and colleagues (7, 12, 44–47). This protocol is exceptionally reliable and as such used by many labs globally. One of the many advantages is that the lack of manual isolation and need for physical removal of retinal tissue from the brain retina hybrid organoids generated using this method. In addition, the protocol does not require the use of 96 well dishes for embryoid body/neurosphere production; while ultra-low attachment 96 well v-bottom plates enable production of uniform embryoid bodies (48), their use greatly reduces throughput, making it difficult to generate retinal organoids in the numbers required for clinical release testing and therapy. By using ReLeSR to lift iPSCs, we demonstrate production of large numbers of embryoid bodies using ultra-low adhesion cell culture flasks. That said, this step requires significant skill, as it is important to not over digest the iPSC culture and obtain cell clusters of uniform size. If large scale production is not required, the 96 well plate approach, which relies on single cell passage, is more easily adopted and has been shown to increase retinal organoid production efficiency (47). Alternatives to 96 well ultra-low adhesion plates for embryoid body production have been developed. These included 6 well AggreWell^TM^ 800 plates, which have 9000 microwells each 800μm in diameter (Stemcell Technologies), and Elplasia^TM^ 12K flasks (Corning), which have 12,000 microwells each 850μm in diameter. As the microwells in these systems are a fraction of the size of a single well of a 96 well plate (e.g., 800μm versus 6700μm), it is yet to be determined if they are useful for production of high-quality retinal organoids.

Unlike embryoid bodies, optic vesicles cannot be generated from single cells using a simple aggregation step. Rather lifting of optic vesicles as intact structures is required. During this process, care should be taken to mechanically dissociate the culture sufficiently to release the optic vesicle from the surrounding tissue, but not so much that the optic vesicles themselves are broken into single cells. For optimal organoid production, manual removal of optic vesicles can be performed (47). As significant skill is required for optic vesicle selection, in the protocol described here, we have opted to lift the entire culture at differentiation day 30 and to select retinal organoids following generation. As retinal organoids have a distinct appearance, their selection is typically easier for technical staff to perform under cGMP.

To demonstrate that the cGMP compliant protocol developed here gives rise to transplantable photoreceptor cells, retinal organoids were dissociated using papain, and liberated cells were injected into the subretinal space of animals that lack photoreceptors at the time transplantation. While non-photoreceptor cells, including RPE, were present within the graft at 3-days post-injection, only human donor photoreceptor cells were identified at 1-month. This likely resulted from the fact that more photoreceptor cells were present in the transplant compared to other cell types (i.e., as per **Figure 5** most human cells present at 3-days post-transplant expressed photoreceptor cell markers). As with previously published protocols, retinal organoids generated using the cGMP compliant retinal differentiation protocol reported here are laminated with photoreceptor cells contained within the outer most layer, separated from the inner retinal neurons by the outer plexiform layer. This organization enables photoreceptor cell enrichment simply by modifying the standard dissociation protocol. Specifically, we have recently demonstrated that partial enzymatic digestion in the absence of physical dissociation liberates photoreceptor cells leaving the inner retinal neurons behind (9). While we performed a complete dissociation here, we did incorporate a filtration step using a 40um cell strainer. This removed any undissociated clusters, which predominantly contain inner retinal neurons and RPE cells, greatly reducing the amount of non-photoreceptor cells present in the transplant. That said, it is possible that only donor photoreceptor cells remained at 1-month because they benefited from trophic support afforded by functional integration with host bipolar cells (**Figure 6**).

As evident in **Figures 5 and 6**, despite aggressive immune suppression, cell survival decreases drastically between 3-days and 1-month following subretinal injection. While it is tempting to attribute poor cell survival to a xenograft response, in 2005 Tomita and colleagues demonstrated that fewer than 10% of transplanted mouse retinal progenitor cells survive to 4-weeks following single cell injection in retinal degenerative mice on the same genetic background (49). Cell survival was increased by approximately 10-fold when donor cells were seeded onto a biodegradable polymer and subsequently transplanted into the subretinal space as an intact graft (49). The drastic difference in donor cell survival seen in this study can likely be attributed in part to sheer stress during injection of single cells using a small-bore needle. In addition, the phenomena of anoikis, which is broadly defined as apoptosis resulting from the loss of cellular adhesion (50), is almost certainly at play. While early polymeric scaffolds greatly enhanced cell survival, fabrication approaches largely resulted in production of planer materials that failed to promote photoreceptor cell polarization and morphologic maturation (51–53). In addition, significant quantities of materials such as poly(lactic-co-glycolic acid) (PLGA) and Poly(glycerol sebacate) (PGS) are not well tolerated in the subretinal space. The lack of biocompatibility of PLGA or PGS-based scaffolds can largely be attributed to the rapid rate of degradation and release of acidic degradation products (54). To address these issues, we have pioneered the use of two-photon lithography to fabricate Polycaprolactone (PCL) based cell delivery scaffolds. Unlike other synthetic polyesters, PCL degrades slowly via hydrolysis of ester linkages (55), which prevents accumulation of acidic degradation products and increases donor cell viability (56). Similarly, two-photon lithography affords the resolution required to fabricate scaffolds designed to promote stacking of donor photoreceptor cells above a monolayer of RPE, creating a graft that more closely mimics the native tissue. We have previously demonstrated that PCL-based cell delivery scaffolds are well tolerated in translational models of cell transplantation (57, 58).

In summary, in this study we report development of a xeno-free 3D retinal differentiation protocol that can be used to produce transplantable photoreceptor cells under cGMP. By removing dispase, Matrigel and FBS from our differentiation protocol, we can be confident that our process is consistent and that the impact of reagent variability on the potency of our clinical product is minimal. Furthermore, we can now generate clinical grade retinal cell grafts for the treatment of patients with advanced retinal degenerative blindness. To translate this technology to the clinic, the next step will be to evaluate safety and efficacy in a large animal model of retinal degeneration.

## Acknowledgments and funding sources

The authors would like to thank the National Eye Institute (EY033331 and EY036143) and the National Science Foundation for providing funding for this work.

## Competing Interests

KAP is a paid consultant and shareholder in Cell X Technologies Inc.

