## Supplemental Data for "Production of clinical grade patient iPSC-derived 3D retinal organoids containing transplantable photoreceptor cells"

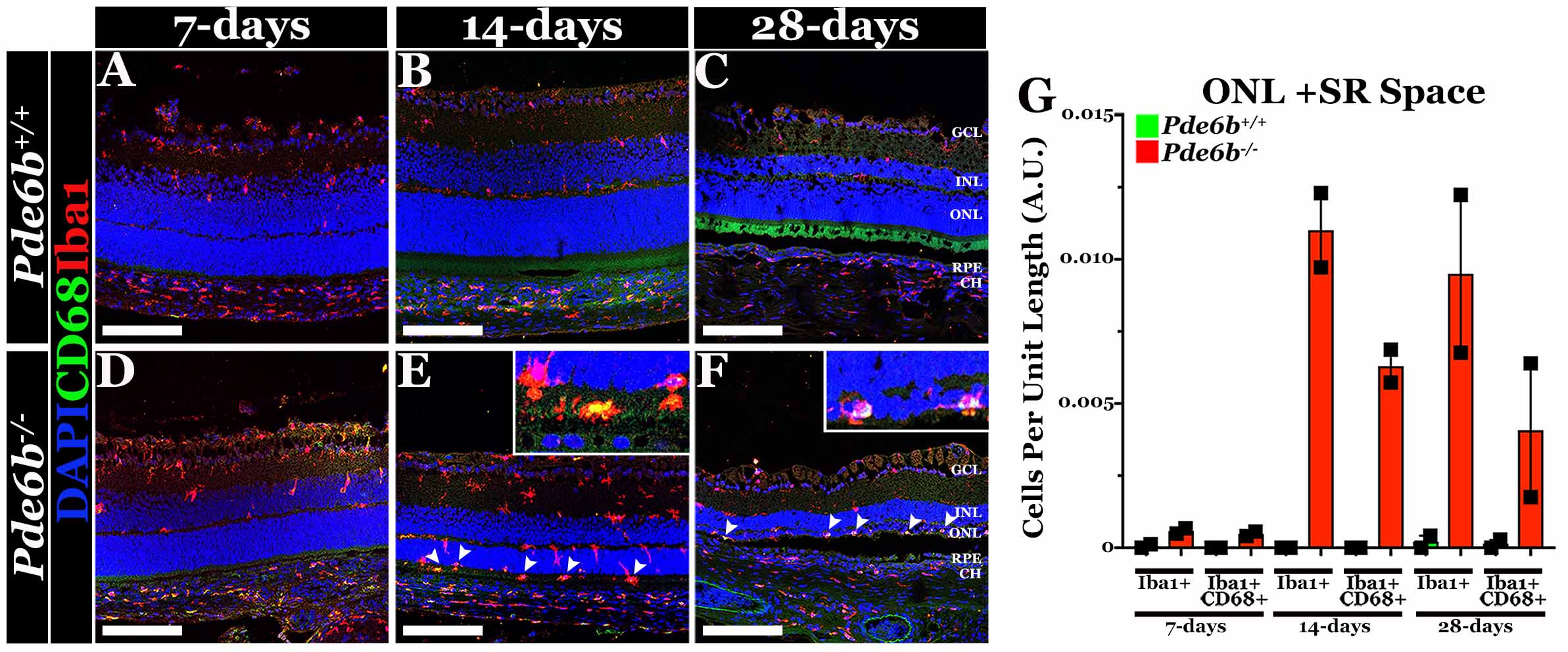

**Supplemental Figure 1. Loss of *Pde6b* causes activation and migration of resident retinal microglia to the outer nuclear layer. A-F)** Representative confocal images labeled with anti-CD68 (green) and anti-Iba1 (red) in unaffected (*Pde6b^+/+^*; A-C) versus knockout (*Pde6b^-/-^*; D-E) at 7-days (A, D), 14-days (B, E) and 28-days (C, F) of age. Cell nuclei are stained with DAPI (blue). Scale bars = 50μm. Abbreviations of retinal layers: Ganglion Cell Layer (GCL), Inner Nuclear Layer (INL), Outer Nuclear Layer (ONL), Retinal Pigmented Epithelium (RPE) and Choroid (CH). **G)** Bar histograms comparing quantification of Iba1-positive (Iba1+) and Iba1/CD68-positive (Iba1+/CD68+) resident retinal microglia per unit length of retina between unaffected (*Pde6b^+/+^*) and knockout (*Pde6b^-/-^*) at 7-, 14- and 28-days of age. Error bars represent standard error of the mean.

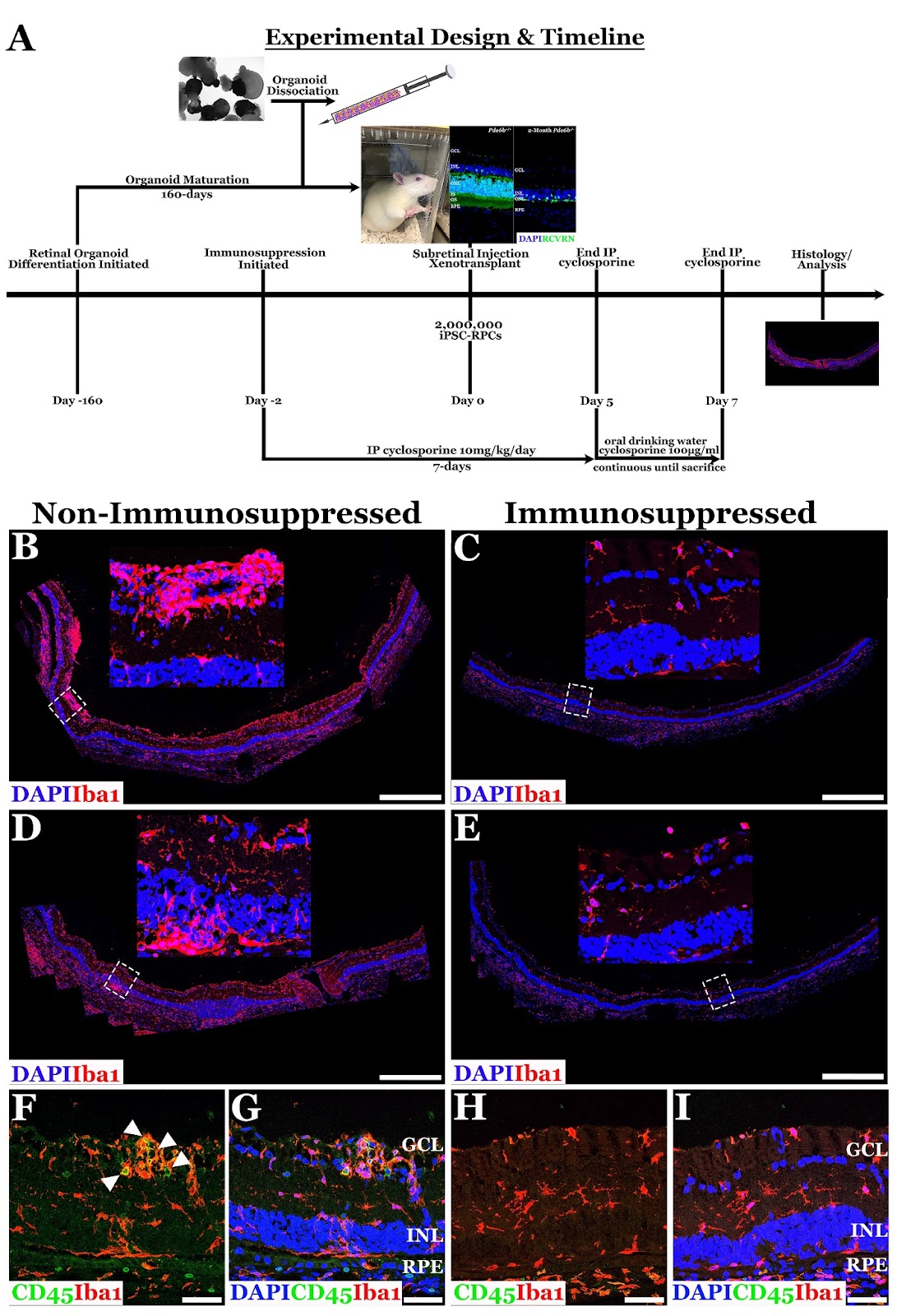

**Supplemental Figure 2.** **The combination of injected and water-supplemented cyclosporine is sufficient for immunosuppression of *Pde6b KO* animals during xenotransplantation of human iPSC-derived retinal cells. A)** Schematic outlining experimental design and timeline for immunosuppression, subretinal injection and analysis of *Pde6b KO* rats. **B-E)** Representative confocal panoramic images labeled with anti-Iba1 (red) to demarcated both resident retinal microglia and infiltrating monocytes in two independently immunosuppressed animals (B, D) and two non-immunosuppressed rats (C, D). Dashed white boxes demarcate areas of higher magnification displayed above each panoramic. **F-I)** Immunofluorescent images of eyes labeled with anti-CD45 (E, G) to label infiltrating monocytes and anti-Iba1 (red). Cell nuclei fluorescently counterstained with DAPI (B-I). Abbreviations for retinal layers in G, I: Ganglion Cell Layer (GCK), Inner Nuclear Layer (INL) and Retinal Pigmented Epithelium (RPE). Scale bars = 500μm (B-E) and 50μm (F-I).

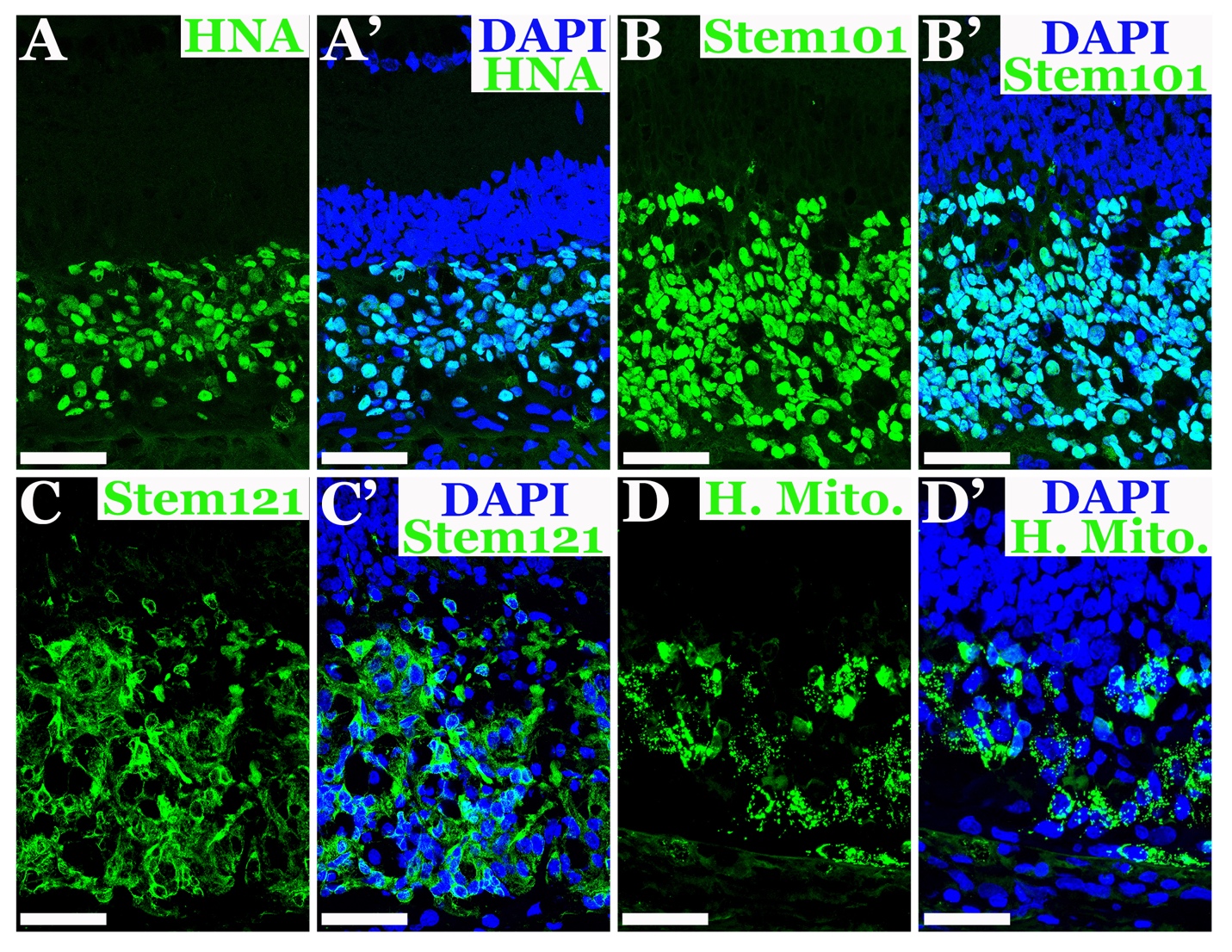

**Supplemental Figure 3. Human iPSC-derived retinal progenitor cells express human-specific markers within the subretinal space of *Pde6b* KO rats at 3-days post-injection. A-D’)** Representative confocal images of human iPSC-derived retinal progenitor cells within the subretinal space of *Pde6b*-deficient rats at 3-days post-injection labeled with anti-human nuclear antigen (HNA; green; A-A’), anti-Stem101 (also known as Ku80; green; B-B’), anti-Stem121 (green; C-C’) and anti-human mitochondria (H. Mito; green; D-D’). Merged images include the nuclear counterstain, DAPI (blue). Scale bars = 50μm.

**Table S1: Antibodies used**

| **Primary Antibodies** | | |
| --- | --- | --- |
| **Antibody** | **Company** | **Catalog Number** |
| OTX2 | R&D Systems | AF1979 |
| Recoverin | MilliporeSigma | AB5585 |
| PKCalpha | Cell Signaling | 2056S |
| NRL | R&D Systems | AF2945 |
| S opsin (blue cone) | MilliporeSigma | AB5407 |
| ARR3 | LifeSpan Bio | LS-C368677 |
| Calretinin | Abcam | ab702 |
| Synaptophysin | Agilent | MO776 |
| CRALBP | Abcam | ab15051 |
| HNA | MilliporeSigma | MAB1281 |
| Stem101 | Takara Bio | 440400 |
| Stem121 | Takara Bio | 440410 |
| Human mitochondria (HMA) | MilliporeSigma | MAB1273 |
| Iba1 | Wako Chemicals | 019-19741 |
| CD68 | Bio-Rad | MCA341GA |
| CD45 | Abcam | ab33923 |
| **Fluorescently-Conjugated Secondary Antibodies** | | |
| **Antibody** | **Company** | **Catalog Number** |
| Goat anti-mouse 488 | Thermo Fisher Scientific | A32723 |
| Goat anti-mouse 555 | Thermo Fisher Scientific | A32727 |
| Goat anti-mouse 647 | Thermo Fisher Scientific | A28181 |
| Goat anti-rabbit 488 | Thermo Fisher Scientific | A32731 |
| Goat anti-rabbit 546 | Thermo Fisher Scientific | A11035 |
| Goat anti-rabbit 647 | Thermo Fisher Scientific | A32733 |
| Donkey anti-goat 546 | Thermo Fisher Scientific | A11056 |
| Donkey anti-goat 647 | Thermo Fisher Scientific | A32849 |
| Donkey anti-goat 488 | Thermo Fisher Scientific | A11055 |
| Donkey anti-rabbit 647 | Thermo Fisher Scientific | A31573 |
